# *TFmapper*: A tool for searching putative factors regulating gene expression using ChIP-seq data

**DOI:** 10.1101/262923

**Authors:** Jianming Zeng, Gang Li

**Affiliations:** Faculty of Health Sciences, University of Macau, Macau, China

**Author notes:** Email: Zeng Jianming, Gang Li.

**Keywords:** Chromatin immunoprecipitation, Next-generation sequencing, Regulation of gene expression

## Abstract

**Background:** Next-generation sequencing coupled to chromatin immunoprecipitation (ChIP-seq), DNase I hypersensitivity (DNase-seq) and the transposase-accessible chromatin assay (ATAC-seq) has generated enormous amounts of data, markedly improved our understanding of the transcriptional and epigenetic control of gene expression. To take advantage of the availability of such datasets and provide clues on what factors, including transcription factors, epigenetic regulators and histone modifications, potentially regulates the expression of a gene of interest, a tool for simultaneous queries of multiple datasets using symbols or genomic coordinates as search terms is needed.

**Results:** In this study, we annotated the peaks of thousands of ChIP-seq datasets generated by ENCODE project, or ChIP-seq/DNase-seq/ATAC-seq datasets deposited in Gene Expression Omnibus and curated by CistromeDB; We built a MySQL database called TFmapper containing the annotations and associated metadata, allowing users without bioinformatics expertise to search across thousands of datasets to identify factors targeting a genomic region/gene of interest in a specified sample through a web interface. Users can also visualize multiple peaks in genome browsers and download the corresponding sequences.

**Conclusion:** TFmapper will help users explore the vast amount of publicly available ChIP-seq/DNase-seq/ATAC-seq data, and perform integrative analyses to understand the regulation of a gene of interest. The web server is freely accessible at http://www.tfmapper.org/.

## Background

Next-generation sequencing (NGS) coupled to chromatin immunoprecipitation (ChIP-seq), DNase I hypersensitivity (DNase-seq) and the transposase-accessible chromatin assay (ATAC-seq) provide powerful tools for studying gene regulation by factors in cis and trans, which includes components of the basal transcriptional machinery, transcription factors, chromatin regulators, histone variants, histone modifications and others. A great amount of data of ChIP-seq, DNase-seq and ATAC-seq has been accumulated by individual laboratories and large-scale collaborative projects, including the ENCODE (Encyclopedia of DNA Elements) Consortium [1], the Roadmap Epigenomics Mapping Consortium (Roadmap) [2] and the International Human Epigenome Consortium (IHEC) projects [3]. The datasets are usually publicly accessible through the Gene Expression Omnibus (GEO) of the National Center for Biotechnology Information (NCBI) [4], and the data portals of large-scale projects.

Despite the easy access, mining and interpreting the ChIP-seq/DNase-seq/ATAC-seq data is challenging for regular users, especially for bench biologists with limited bioinformatic expertise. Adding to the complexity, the data qualities and data analysis pipelines are remarkably varied, which hinder their direct use in further analyses [5]. To address the challenges, multiple algorithms/metrics have been developed to evaluate the data quality bioinformatically, such as ENCODE quality metrics, NGS-QC, ChiLin and others [6-9]. NGS-QC developed by Gronemeyer and colleagues built a quality control (QC) indicator database of the largest collection of publicly available NGS datasets [6, 8], which provides a solid start point for further analysis. CistromeDB developed by Liu and colleagues curated and processed a huge collection of human and mouse ChIP-seq and chromatin accessibility datasets from GEO with a standard analysis pipeline ChiLin [9], and further evaluated individual data quality under several scoring metrics [10]. High-quality processed ChIP-seq data generated by ENCODE consortium, including histone modification, chromatin regulator and transcription factor binding data in a selected set of biological samples, are also available through its data portal [11]. ENCODE and CistromeDB provide access to the processed data, and the corresponding metadata including the sources and properties of biological samples, experimental protocols, the antibody used and others, which offer opportunities for users to re-analyze the data and identify the genome-wide targets of a transcription regulator in different cell lines and tissues. Nonetheless, different questions are often asked, such as: Which transcription factors are responsible for the regulation of a gene of interest, and what is the epigenetic landscape of a gene of interest in a particular tissue or cell type? To address these questions, a tool for simultaneous queries of multiple datasets using symbols or genomic coordinates of target genes as search terms is needed.

In this study, we collected and annotated a large number of ChIP-seq, DNase-seq and ATAC-seq datasets including the ENCODE datasets and the GEO datasets curated by CistromeDB [10]. We built a Structured Query Language (SQL) database containing the annotations and the associated metadata, allowing users to search across multiple datasets and identify the putative factors which target a specified gene or genomic locus based on actual NGS data. We also provide links for users to visualize the peaks and download the corresponding sequences. In addition, we included an example to demonstrate the utility of *TFmapper*.

## Construction and content

### Data collection

We downloaded the GEO datasets processed by Liu and colleagues from CistromeDB as BED (Browser Extensible Data) files, which includes 6092 ChIP-seq datasets for trans-acting factors and 6068 ChIP-seq datasets for histone modifications generated from human samples; 4786 ChIP-seq datasets for trans-acting factors and 5002 ChIP-seq datasets for histone modifications generated from mouse samples; and 1371 DNase-seq/ATAC-seq datasets. For ENCODE datasets, we downloaded the conservative IDR (Irreproducible Discovery Rate)-thresholded peaks, which are called after combinational analysis of two replicates for each ChIP-seq experiment. The ENCODE datasets selected in this study include 955 transcription factor and 771 histone modification datasets for Homo sapiens, and 145 transcription factor and 1252 histone modification datasets for Mus musculus.

### Data processing

We used HOMER [12] to annotate the peaks in BED files per the newest reference genome, GRCh38/hg38 for human and GRCm38/mm10 for mouse respectively. HOMER assigns each peak to the nearest gene by calculating the distance between the middle of a peak and the transcription start site (TSS) of a gene. Meanwhile, seven genomic features based on RefSeq annotations were assigned to each peak, which are promoter (by default defined from −1kb to +100bp of TSS), TTS region (transcription termination site, by default defined from −100 bp to +1kb of TTS), Exon, 5’ UTR Exon, 3’ UTR Exon, Intron and Intergenic.

### Database design

To enhance the searching speed in identifying all the peaks targeting a selected gene or specific region, we split the table into subsets based on species, sources, factors, and chromosomes, resulting in 176 tables containing the peaks information. We organized the gene location information according to the longest annotation of the transcript in the Consensus CDS (CCDS) database [13], and only the genes which have peaks in any of the ChIP-seq datasets will be kept for searching. The metadata for GEO and ENCODE datasets were re-organized and uploaded to a MySQL database. The R package GEOmetadb [14] is also used to retrieve further annotation details of the samples in GEO.

### Web application

The web application was generated by the R package Shiny and hosted on a Linux server (Ubuntu 16.04), can be accessed by a web browser at http://www.tfmapper.org/. The page layout was made using the R package Shinydashboard, and the interactive tables were displayed using the R package DT.

### Utility and discussion

The primary purpose of *TFmapper* is to search all experimental ChIP-seq datasets and identify the trans-acting factors or histone modifications which show peaks at a gene of interest or a specified genomic region in a user-defined biological sample. To achieve this objective, we downloaded ENCODE and GEO/CistromeDB data, and annotated the peaks using HOMER software [12] based on the peaks called by ENCODE and CistromeDB. We then built a MySQL database called TFmapper containing the annotations and associated metadata with a web interface (Fig. 1), which allows searching of thousands of GEO/ENCODE datasets simultaneously to find what TF/histone marks target a gene of interest in a user-specified biological sample. Currently, TFmapper includes 26445 human and mouse datasets, including 11978 ChIP-seq datasets for trans-acting factors, 11073 ChIP-seq datasets for histone marks/variants, and 1371 DNase-Seq/ATAC-seq datasets (Fig. 1).

**Figure 1.**
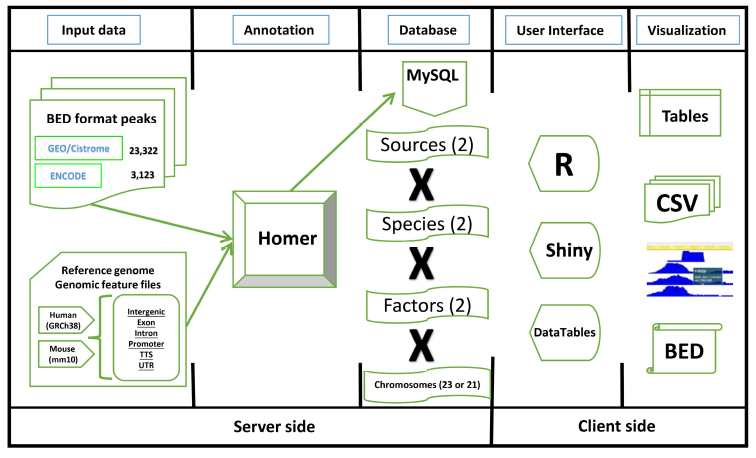
The workflow diagram of *TFmapper*. Peak files in the BED (Browser Extensible Data) format are downloaded from the GEO/Cistrome and the ENCODE data portals. Peaks are annotated to genomic features (Promoter, TTS, 5‘UTR, 3‘UTR, Intron, Exon, Intergenic) using the software HOMER with GRCh38/hg38 (Human) and GRCm38/mm10 (Mouse) as the reference genomes. The annotation results are stored in a MySQL database. To increase the speed of query processing, the peaks are split by species, sources, factors, and chromosomes. For the client side, all the elements on the HTML page are built by R (Shiny), and result tables are created with the DataTables JavaScript library. Results can be downloaded in the CSV or BED format, and peaks can be directly visualized in the WashU Epigenome Browser or the UCSC Genome Browser.

### Website interface

The ‘home’ page of *TFmapper* consists of three sections (Fig. 2).

**Figure 2.**
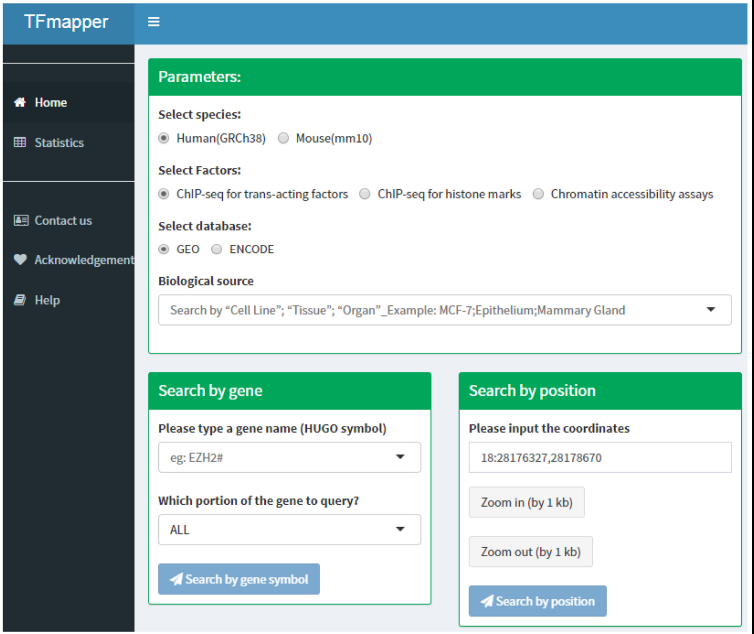
The homepage of *TFmapper*. The Parameters and Search sections are shown here.

#### Parameters Section

In this section, users can define the searching criteria, including species (human or mouse), types of experiments (ChIP-seq for trans-acting factors, ChIP-seq for histone marks or Chromatin accessibility assays), sources of datasets (GEO or ENCODE) and biological sources. Each biological source is defined by three properties. Based on the original description, the properties could be ‘cell line’, ‘tissue’ and ‘organ’. All three properties are searchable.

#### Search Section

Users can query the database by 1) gene symbols or 2) genomic coordinates of GRCh38/hg38 (Homo Sapiens) or GRCm38/mm10 (Mus Musculus) respectively. Users can further define which portion of the gene to query (Fig. 2).

#### Results Section

The search results are displayed in the form of a table including the following fields: Sample identifiers (GEO sample (GSM) IDs or ENCODE IDs); SampleID defined by CistromeDB; name of factors (trans-acting factors, histone modifications); links for peak visualization in the WashU Epigenome Browser [15] or the UCSC Genome Browser [16]; the coordinates of the peak; distance between the center of a peak and the transcription start site (TSS) of a gene; the enrichment scores, the p- and q- values for ChIP-seq peaks called by Model-based Analysis for ChIP-Seq (MACS2) [17, 18]; the genomic attribute (promoters, intergenic or intragenic peaks); title and source name of the dataset, etc. At the bottom of this section, a graph is automatically generated to display the number of experiments being performed for each factor. The search results are sorted by default per the distance between the feature site and the TSS. All fields can be further sorted by clicking the arrow at the left of the title of each field, or narrowed by specifying the desired value for each field. Under the ‘distance’ field, a slider will show up when clicked; the user can move the slider to narrow down the region. Several other fields are clickable; users will be redirected to the specific GSM or ENCODE information page by clicking the sample identifier field, and the user can retrieve the actual sequence of a peak by clicking the peak coordinates. The query results are stored in a CSV (Comma Separated Values) format file, or a BED (Browser Extensible Data) format file. The CSV format file contains all the fields in the search results, and the BED format file contains the names of factors, coordinates and enrichment scores for all peaks. Both files are downloadable for downstream analysis. For example, the user can load the BED format file into the Integrative Genomics Viewer (IGV) to visualize each peak at a specific genomic region [19]. More importantly, a link will appear on the top of the table when multiple peaks are highlighted in blue, which allows users to visualize multiple peaks in the WashU Epigenome Browser (Fig. 3).

**Figure 3.**
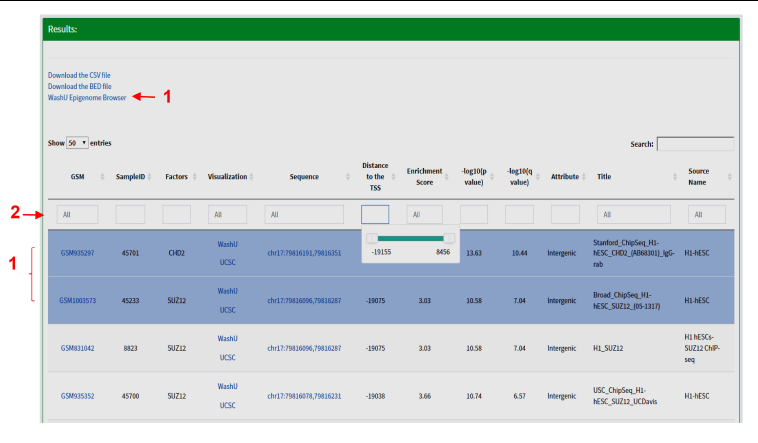
The ‘Results’ section of *TFmapper*. When multiple peaks are selected, a link for visualization in the WashU Epigenome Browser will appear (red arrow 1). The results can be further filtered by typing search terms in the boxes (red arrow 2); a slider will appear when the box under ‘Distance’ being clicked, which can be moved to narrow down the region.

### Case studies

To demonstrate the utility of *TFmapper*, we examined the transcriptional regulation of chromobox homologs (CBXs) in the H1 line of the human embryonic stem cells (H1-hESC). CBX proteins (including CBX2, CBX4, CBX6, CBX7, and CBX8) are mammalian homologs of Drosophila Polycomb protein, bind to trimethylated lysine 27 of histone H3 (H3K27me3) through the chromodomain. Orthologs of Cbxs are differentially expressed in mouse embryonic stem cells, in which Cbx7 is highly expressed. Cbx7 represses the expression of other CBXs and contributes to the maintenance of stem cell pluripotency [20, 21]. Nonetheless, analysis of RNA-Seq data of ENCODE clearly demonstrated that the expression profiles of CBXs are strikingly different in human ESCs than in mouse ESCs (Fig.4a). In H1 cells, CBX2 are highly expressed, CBX6 and CBX7 are moderately expressed, whereas CBX4 and CBX8 are lowly or not expressed. Human CBX2, CBX8 and CBX4 are sequentially localized on chromatin 17. To understand why CBX2 and its neighbor CBX8 express differently in H1 cells, we used *TFmapper* to retrieve information on chromatin regulators that target CBX2 and CBX8 (Supplementary Table S1, S2), and the statuses of histone modifications on CBX2 and CBX8 gene in H1 cells.

To our surprise, although CBX2 and CBX8 are expressed at significantly different levels, the factors targeting CBX2 and CBX8 are largely overlapped, and include those correlated with either gene activation or silencing (Fig. 4b). However, there are several factors which target CBX8 only, such as BRCA1, CHD7, CTBP2, MAFK, MXI1, POU5F1, SOX2, SP2, SRF and USF2 (Fig. 4b, Supplementary Fig. S1). Only GTF2F1, which encodes the alpha subunit of the general transcription factor TFIIF and promotes transcription elongation, targets CBX2 exclusively (Supplementary Fig. S1). As shown in Figure 4c, a CTCF peak sits proximally upstream (-500bp) of CBX2 transcription start site, a double peak of CTCF present in the intergenic region between CBX2 and CBX8 in H1 cells, suggesting CBX2 and CBX8 sit in two neighborhoods, possibly insulated by the intergenic CTCF double peak. A wide binding of EZH2 and enrichment of repressive mark H3K27me3 were found on CBX8, extending almost continuously from upstream 19 Kb of CBX8 transcription start site (TSS) to downstream 2 Kb of CBX8 transcription termination site (TTS), suggesting CBX8 is silenced by Polycomb Repressive Complex 2 (PRC2).

**Figure 4.**
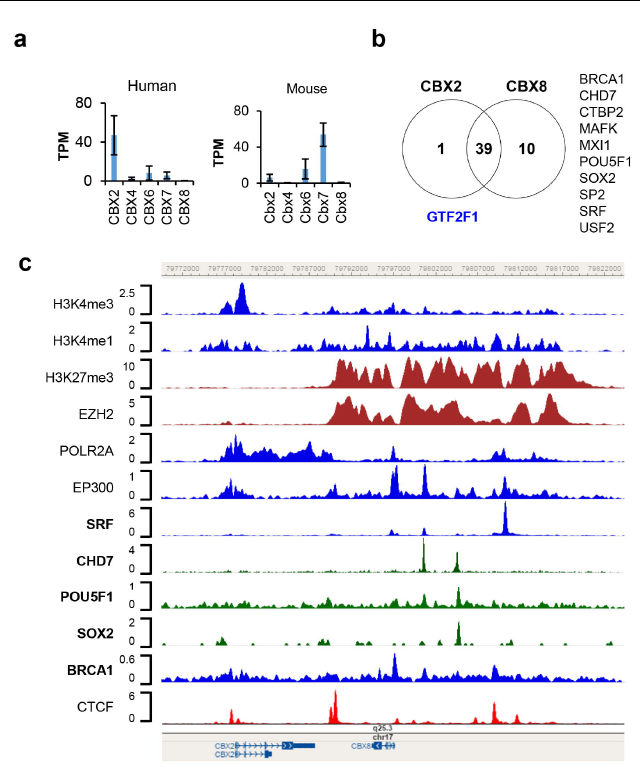
Transcriptional regulation of CBX2 and CBX8 in hESC-H1 cells. (**a**) mRNA expression of CBX2, CBX4, CBX6, CBX7 and CBX8 in human and mouse embryonic stem cells (hESCs and mESCs). The data are retrieved from the RNA-seq datasets of ENCODE. The expression values are presented as transcripts per million (TPM). (**b**) Venn diagram of trans-acting factors targeting CBX2 and CBX8 in hESC-H1 cells. The detailed information on the factors can be found in the Supplementary Table S1 and S2. The factors targeting CBX2 or CBX8 exclusively are indicated in blue and black respectively. (**c**) Binding patterns of selected trans-acting factors and histone modification marks at the CBX2-CBX8 locus in hESC-H1 cells. The peaks are visualized with the WashU Epigenome Browser. The GEO accession number for each dataset can be found in the Supplementary Table S3.

Further detailed examination of the trans-acting factors targeting CBX8 reveals several other intriguing features which are worth to be further investigated: 1) a putative poised enhancer region centered at the serum response factor (SRF) peak, 13Kb upstream of the TSS of CBX8. Around the SRF peak, there are significant POLR2A binding, EP300 binding, H3K27me3 and H3K4me1 enrichment, strongly suggests the existence of a poised enhancer, and SRF might regulate CBX8 expression. 2) a BRCA1 peak at the TSS of CBX8. A significant BRCA1 peak was found at the TSS of CBX8, suggesting the potential involvement of BRCA1 in the regulation of CBX8 expression. Recently, Oza J et al. reported that CBX8 is quickly recruited to the sites of DNA damage [22]. The targeting of CBX8 by BRCA1 further supports the participation of CBX8 in DNA damage response. 3) POU5F1 (OCT4), SOX2 and CHD7 target CBX8. Two significant CHD7 peaks were found 3.5 Kb and 7.5 Kb upstream of CBX8 TSS site; the later peak overlaps with POU5F1 and SOX2 peaks. The transcriptional circuitry mediated by OCT-4 and SOX2 supports the self-renewal and pluripotency of ESC. OCT-4 and SOX2 work cooperatively to regulate genes primarily as transcriptional activators [23]. The chromodomain helicase DNA-binding protein 7 (Chd7) physically interacts Sox2 [24], targets active gene enhancer [25], and maintains open chromatin configuration [26, 27]. The peaks of POU5F1, SOX2, and CHD7 on the distal promoter region of CBX8 partly overlap, or reside in between the broad and strong H3K27me3 peaks. We hypothesize these factors might work together to prime CBX8 to be expressed. Whether the hypothesis holds true requires further investigation.

A large amount of ChIP-seq, DNase-seq and ATAC-seq data is freely available in the public domain. Unfortunately, the majority of the datasets deposited in GEO of NCBI are not processed, which poses a significant barrier to datamining by non-bioinformaticians. CistromeDB reprocessed a huge collection of human and mouse ChIP-seq and chromatin accessibility datasets from GEO with a standard analysis pipeline, and calculated metrics reflecting data quality, in turn, CistromeDB provides a very useful resource for datamining. However, Cistrome only allows factor-oriented (in trans) queries: to identify the target sites of a specific transcription factor or histone mark on the genome in a specified sample, but not gene-oriented (in cis) query: to identify all of the factors targeting a gene of interest in a specific biological sample by searching multiple datasets. To address this problem, we developed TFmapper. To our knowledge, other tools including BindDB [28], GTRD [29] and Remap [30] are capable of performing similar functions as TFmapper. However, BindDB only includes 455 datasets generated from mouse and human ESCs and iPSCs. The query result from BindDB does not include the exact peak coordinates, making it difficult to analyze the relationships between factors. It does not provide adequate descriptive information on the samples and clickable links to GEO. In addition, BindDB does not support sequence retrieving and peak visualization. Remap includes 1085 human datasets of transcription factors, while GTRD includes 8828 datasets of transcription factors for human and mouse, both tools do not contain datasets for histone marks and chromatin accessibility assays. Both tools merged peaks from different cell lines with distinct gene expression profiles to facilitate the identification of putative transcription factor binding sites (TFBS) or cis-acting regulatory sequences in the genome. Different types of cells have distinct gene expression profiles: correspondingly, different types of cells might use different enhancers to regulate the same gene and have different binding patterns of transcription factors [31, 32]. Although the merging of peaks serves its intended purpose well, it prevents the users from predicting if a targeting event happens in the biological sample they are studying. In addition, both tools don‘t allow users to trace back to the original GSM dataset easily to check in detail the culture/treatment condition and quality of the data.

## Conclusions

*TFmapper* enables users to search across thousands of ChIP-seq/DNase-seq/ATAC-seq datasets, including the ENCODE datasets and the GEO datasets curated by Cistrome, to identify the putative factors which target a genomic region/gene of interest. We believe this tool will help eliminate bioinformatic barriers, allow the scientists to take advantage of the enormous amount of existing ChIP-seq data and perform integrative analysis to understand mechanisms of gene regulation.

## List of abbreviations

NGS: Next-generation sequencing
GEO: the Gene Expression Omnibus
NCBI: the National Center for Biotechnology Information
IHEC: the International Human Epigenome Consortium
ENCODE: the Encyclopedia of DNA Elements) Consortium
ChIP-Seq: Next-generation sequencing coupled to chromatin immunoprecipitation
DNase-seq: Next-generation sequencing coupled to DNase I hypersensitivity
ATAC-seq: Next-generation sequencing coupled to the transposase-accessible chromatin assay.

## Declarations

### Ethics approval and consent to participate

No ethics approval was required for the study.

### Consent for publication

Not applicable.

### Availability of data and material

TFmapper is freely accessible at http://www.tfmapper.org/. The complete source code for TFmapper is freely available at GITHUB (https://github.com/jmzeng1314/TF_map) under a GPLv3 license.

### Authors’ contributions

J.Z. and G.L. conceived the study. J.Z. and G.L. collected and analyzed the data. J.Z. built the database and the Web server. All authors wrote and approved the manuscript.

## Acknowledgements

Not applicable.

